# Prediction of non-canonical routes for SARS-CoV-2 infection in human placenta cells

**DOI:** 10.1101/2020.06.12.148411

**Authors:** F. B. Constantino, S. S. Cury, C. R. Nogueira, R. F. Carvalho, L. A. Justulin

**Author notes:** These authors contributed equally (F. B. Constantino, S. S. Cury). **Corresponding authors**, Corresponding author’s information: Luis Antonio Justulin, Department of Structural and Functional Biology, Institute of Biosciences, São Paulo State University (UNESP), CEP: 18.618-689, Botucatu, São Paulo – Brazil, Robson Francisco Carvalho, Department of Structural and Functional Biology, Institute of Biosciences, São Paulo State University (UNESP), CEP: 18.618-689, Botucatu, São Paulo – Brazil.

## Abstract

The SARS-CoV-2 is the causative agent of the COVID-19 pandemic. The data available about COVID-19 during pregnancy have demonstrated placental infection; however, the intrauterine transmission of SARS-CoV-2 is still debated. Intriguingly, while canonical SARS-CoV-2 cell entry mediators are expressed at low levels in placental cells, the receptors for viruses that cause congenital infections such as the cytomegalovirus and Zika virus are highly expressed in these cells. Here we analyzed the transcriptional profile (microarray and single-cell RNA-Seq) of proteins potentially interacting with coronaviruses to identify non-canonical mediators of SARS-CoV-2 infection and replication in the placenta. We show that, despite low levels of the canonical cell entry mediators *ACE2* and *TMPRSS2*, cells of the syncytiotrophoblast, villous cytotrophoblast, and extravillous trophoblast co-express high levels of the potential non-canonical cell-entry mediators *DPP4* and *CTSL*. We also found changes in the expression of *DAAM1* and *PAICS* genes during pregnancy, which are translated into proteins also predicted to interact with coronaviruses proteins. These results provide new insight into the interaction between SARS-CoV-2 and host proteins that may act as non-canonical routes for SARS-CoV-2 infection and replication in the placenta cells.

## Introduction

The severe acute respiratory syndrome coronavirus 2 (SARS-CoV-2) is the causative agent of the coronavirus disease 2019 (COVID-19) (*1*). It was first notified at the end of 2019, in Wuhan, China, and become a worldwide pandemic (*2*). At the beginning of July, six months later, COVID-19 infected over 12 million people and is the cause of approximately 557,000 deaths worldwide (https://coronavirus.jhu.edu/).

Older age, laboratory abnormalities, and several comorbidities are associated with the more severe cases of COVID-19 (*3*). For specific groups of COVID-19 patients, for example, pregnant women, the potential impacts of SARS-CoV-2 infection remains mostly unknown, and data are limited. However, considering previous works reporting coronaviruses infections (*4*), pregnant women are at higher risk of SARS-CoV-2 infection due to physiological changes in the immune, cardiorespiratory, and metabolic systems (*5*).

Although only a small number of maternal viruses infections are transmitted to the fetus, some may cause life-threatening diseases (*6*). These viruses use cellular host entry mediators expressed by placenta cells, as described for the cytomegalovirus and the Zika virus (*7*, *8*), to infect these cells. The Zika virus (ZIKV) outbreak, associated with fetal brain damage, emphasizes the necessity of further characterization and understanding placental infection or intrauterine (vertical) transmission of SARS-CoV-2, as well as the possible adverse fetal outcomes. The few studies on the subject have provided contradictory findings, with some reports suggesting no evidence of placental infection or vertical transmission of SARS-CoV-2 (*9*–*11*). Conversely, multiple lines of evidence have shown placental SARS-CoV-2 infection in pregnant women diagnosed with moderate to severe COVID-19. Neonates born from mothers with COVID-19 presented a positive serological test for SARS-CoV-2 immunoglobulin (Ig)M and IgG (*12*, *13*). While the IgG can be transferred from mother to fetus across the placenta, the detection of IgM in newborns suggests a vertical transmission of the virus, since IgM cannot cross the placental barrier due to its high molecular mass (*14*). Accordingly, SARS-CoV-2 RNA transmission was comprehensively confirmed by pathological and virological investigations. (*15*) Also, it was recently shown SARS-CoV-2 particles in syncytiotrophoblast with generalized inflammation, diffuse perivillous fibrin depositions, and tissue damage in an asymptomatic woman (*16*). Remarkably, these placental alterations due to the SARS-CoV-2 infection lead to fetal distress and neonatal multi-organ failure. These results highlight the importance of exploring the expression profile of potential host mediators of the SARS-CoV-2 that may create a permissive microenvironment to placental infection and possible vertical transmission of the virus.

In fact, like other viruses, SARS-CoV-2 requires diverse host cellular factors for infection and replication. The angiotensin-converting enzyme 2 (ACE2) is the canonical receptor for the SARS-CoV-2 spike protein receptor-binding domain (RBD) for viral attachment (*17*). This process is followed by S protein priming by cellular transmembrane serine protease 2 (TMPRSS2) that allows the fusion of the virus with host cellular membranes (*17*). Single-cell RNA sequencing (scRNA-Seq) has demonstrated that both *ACE2* and *TMPRSS2* are co-expressed in multiple tissues affected by COVID-19, including airway epithelial cells, cornea, digestive and urogenital systems (*18*). Few cells express *ACE2* and *TMPRSS2* in the placenta (*18*, *19*), suggesting that SARS-CoV-2 is unlikely to infect the placenta through the canonical cell entry mediators. Therefore, other host interacting proteins may play a role in the biological cycle of the virus and contribute to the pathogenesis of SARS-CoV-2 in the placenta. In this paper, we demonstrate, through transcriptomic (microarray and scRNA-Seq) analysis and *in silico* predictions of virus-host protein-protein interactions, that cells of the syncytiotrophoblast, villous cytotrophoblast, and extravillous trophoblast express high levels of potential non-canonical cell-entry mediators dipeptidyl peptidase 4 (*DPP4*) and cathepsin L (*CTSL*), despite low-levels of *ACE2* and *TMPRSS2*. These results open new avenues of investigation of the human placenta infection by the SARS-CoV-2.

## Results

### Prediction of non-canonical routes for SARS-CoV-2 infection in human placenta cells

We first investigated the gene expression in placental tissues of classical host-virus interacting proteins described in the literature. The canonical entry receptors *ACE2* and *TMPRSS2* were low expressed throughout gestation. *CTSL*, which is translated into a lysosomal cysteine proteinase that plays a role in intracellular protein catabolism - presented the highest level of expression in the placenta during the first, second, and third trimester. Similarly, *DPP4*, which is translated into an intrinsic membrane glycoprotein, was highly expressed throughout gestation (**Figure 1A**).

**Figure 1.**
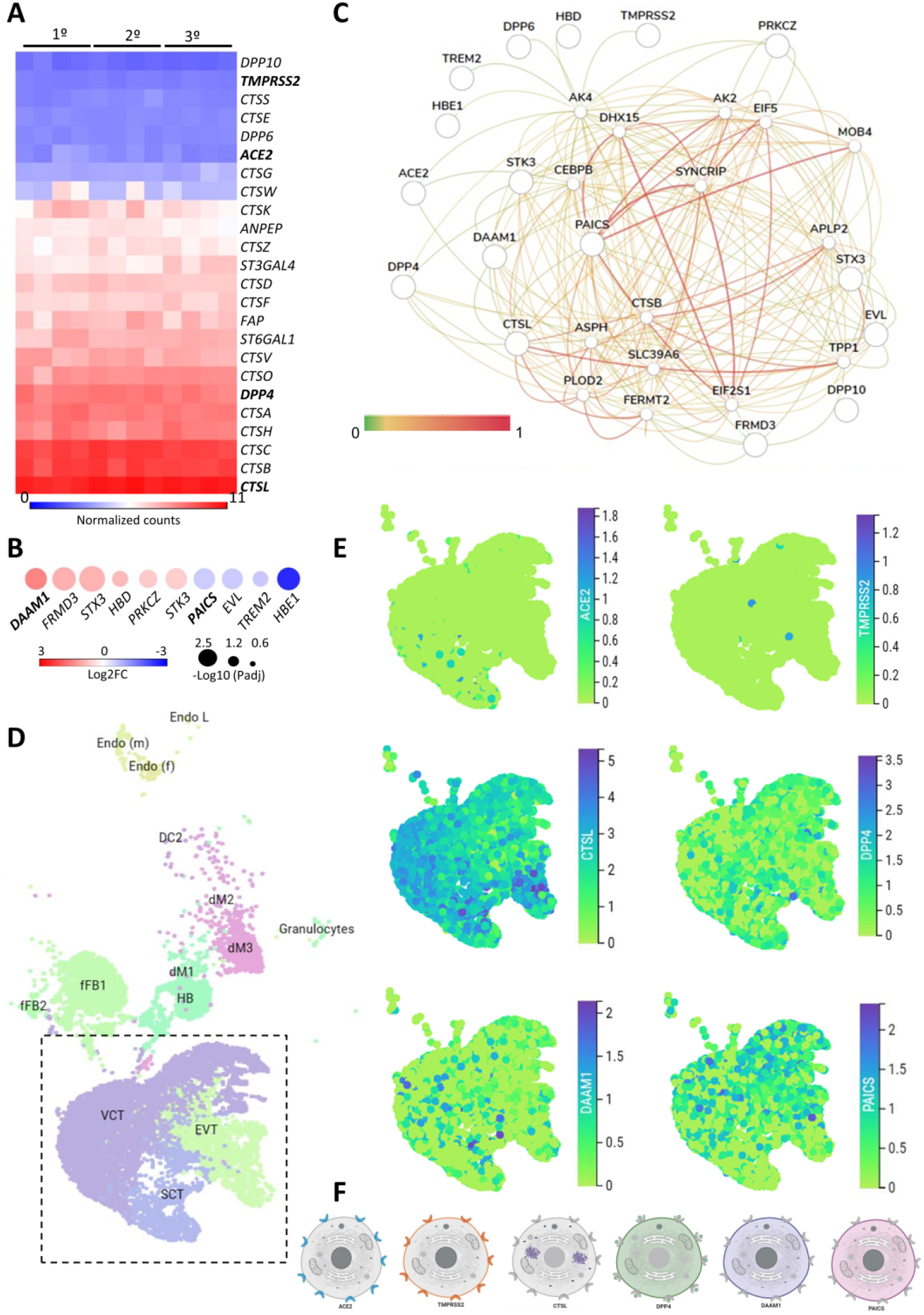
The expression landscape of human placenta proteins potentially interacting with SARS-CoV-2. **A**. Heatmap illustrating gene expression (counts RMA normalized) of SARS-CoV-2 interacting proteins from placental samples in the first, second and third trimester of gestation (GSE9984)(*36*). **B**. Heat-scatter plot presenting ten differentially expressed genes in the placenta that encode proteins potentially interacting with SARS-CoV as predicted by the P-HIPSter (http://phipster.org/) database. The color and size of the circles correspond to the log2FC and −log10 transformed FDR adjusted P-value, respectively. Fold change represents the gene expression of placenta samples from the third trimester of gestation compared with the first trimester. Bolded genes represent the six candidates selected for the further scRNA-seq evaluations. **C**. Tissue-specific gene network of placenta proteins predicted to interact with coronaviruses. The network was generated using the HumanBase (https://hb.flatironinstitute.org/) online tool. **D**. Dimensionality reduction demonstrates different cell populations identified in five first-trimester human placental samples in a Uniform Manifold Approximation and Projection (UMAP) plot, by using scRNA-Seq data from Vento-Tormo et al., (*40*). The images were generated using the COVID-19 Cell Atlas (https://www.covid19cellatlas.org) functionalities. **E**. UMAP plots representing the gene expression (log-transformed normalized counts) of *ACE2*, *TMPRSS2*, *CTSL*, *DPP4*, *DAAM1*, and *PAICS* in human placental cells from the syncytiotrophoblast, villous cytotrophoblast, and extravillous trophoblast. The images were generated using COVID-19 Cell Atlas (https://www.covid19cellatlas.org) functionalities. **F**. Cell illustrations depicting the protein localization of ACE2, TMPRSS2, CTSL, DPP4, DAAM1, and PAICS in cell compartments, as described in the Human Protein Atlas database (http://www.proteinatlas.org). Illustrations were created using BioRender App (https://biorender.com/). RMA: Robust Multichip Average; FC: Fold Change; FDR: false discovery rate; Padj: adjusted P-value; SCT, syncytiotrophoblast; VCT, villous cytotrophoblast; EVT, extravillous trophoblast; HB, Hofbauer cells; FB: fibroblasts; dM, decidual macrophages; Endo, endothelial cells; l, lymphatic; m, maternal; f, fetal.

Next, we analyzed the gene expression profile during gestation in placental tissues. We found 25 differentially expressed genes (DEGs) in the second trimester, and 687 DEGs in the third, when they were independently compared to the first trimester (**Supplementary Table 1**). All DEGs were divided into three clusters by the K-means clustering analysis (**Supplementary Figure 1A**). Cluster 1 includes genes that increase expression during pregnancy, and these genes enriched terms related to blood vessels morphogenesis, complement cascade, extracellular matrix organization, and cellular response to nitrogen compounds. Cluster 2 encompasses genes that increase expression specifically in the third trimesters, which are related to female pregnancy, growth hormone signaling pathway, homeostasis, and steroid biosynthetic process. Cluster 3 includes genes that decrease expression during pregnancy. and these genes enriched terms related to cell division, sulfur compounds biosynthetic process, chromosomal segregation, PID MYC active pathway (**Supplementary Figure 1B and Supplementary Table 2**). The gene expression changes in the course of pregnancy revealed that the second and third trimesters enriched genes that are associated with the placental growth, mainly in the third trimester (**Supplementary Figure 1C and Supplementary Table 3**). Even with a high discrepancy between the number of DEGs in second and third trimesters (25 and 687, respectively), we identified similarities in the enriched terms between gene clusters (**Supplementary Figure 1D**) and between the second and third trimesters of gestation (**Supplementary Figure 1E**).

We further asked whether the DEGs in the placenta during gestation are translated into proteins that potentially interact with SARS-CoV proteins. Considering that the current cases of SARS-CoV-2 placental infection are mainly related to the third trimester of pregnancy (*15*, *20*), we selected the 687 DEGs in the placenta in the third trimester compared to the first trimester and, among them, 474 and 213 were up- and down-regulated, respectively (**Supplementary Table 1**). Next, we selected the human proteins potentially interacting with SARS-CoV using the P-HIPSter database. We found 32 virushost interacting proteins of SARS-CoV (**Supplementary Table 4**), and nine virus-host interacting proteins of the ZIKV (**Supplementary Table 5**). We also found that, from these list of virus-host interacting proteins, 10 DEGs (*DAAM1, FRMD3, STX3, HBD, PRKCZ, CTK3, PAICS, EVL, TREM2*, and *HBE1*) are translated into proteins that interact with SARS-CoV proteins, and one with the ZIKV (**Supplementary Figures 1F and 1G**). Among these 10 DEGs, six were up-regulated (*DAAM1, FRMD3, STX3, HBD, PRKCZ*, and *CTK3*) and four (*PAICS, EVL, TREM2*, and *HBE1*) were down-regulated in the third trimester (**Figure 1B**). The gene *DAAM1*, which is translated into an intrinsic membrane glycoprotein implicated in cell motility, adhesion, cytokinesis, and cell polarity - showed the highest level of fold change in the third trimester of gestation compared to the first (logFC= 1.43; **Figure 1B**). The host-virus protein-protein interactions (PPI) predicted for these 10 DEGs presented a Likelihood Ratio (LR) > 100 (**Supplementary Table 6**), according to P-HIPSTer (*21*). Noteworthy, *PAICS* transcript, which is translated into an enzyme that catalyzes the sixth and seventh steps of the novo purine biosynthesis, were predicted to interact with both SARS-CoV and ZIKV (**Figure 1B and Supplementary Table 7**). We observed a significant interaction of placental proteins predicted to interact with SARS-CoV-2 (based on the DEGs or not), and PAICS was also predicted to interact with other proteins in the PPI network with the highest degree (**Figure 1C and Supplementary Table 8**).

We next used single-cell RNA sequencing to analyze the expression of the 10 DEGs translated into proteins that potentially interact with SARS-CoV in human placental cells (**Supplementary Figures 1H and 1I**). We selected the genes *ACE2, TMPRSS2, CTSL, DPP4, PAICS*, and *DAAM1* for further investigations using scRNA-seq transcriptome data from cells of the syncytiotrophoblast (n=1144), villous cytotrophoblast (n=8244), and extravillous trophoblast (n=2170) of non-disease human placental tissues, considering the potential relevance of these genes for SARS-CoV infection and replication in the organ (**Figure 1D**). We noticed that the expression of the classical SARS-CoV-2 entry receptor genes (*ACE2* and *TMPRSS2*) was minimally expressed in these cells. In contrast, potential non-canonical cell entry mediator genes *DPP4* and *CTSL*, as well as the genes for the predicted virus-host interaction proteins *DAAM1*, and *PAICS* were expressed at higher levels (**Figure 1E and Supplementary Figure 2A**). We also looked for the co-expression of *ACE2* and *TMPRSS2* with *DPP4, CTSL, DAAM1*, and *PAICS* by using scRNA-Seq in these same cells (**Figure 2**). This analysis revealed that *DPP4, CTSL, DAAM1*, and *PAICS* are co-expressed at high-levels, while these genes are co-express at low levels with *ACE2* and *TMPRSS2* (**Figure 2**). Remarkably, we found only three cells co-expressing *ACE2* and *TMPRSS2*. For this reason, we consider that *DPP4, CTSL, DAAM1*, and *PAICS* may represent candidates of an alternative route for SARS-CoV-2 infection in human placentas. We used The *Human Protein Atlas* to predict the subcellular location of proteins encoded by our candidates. DPP4 is predicted as intracellular, membrane, and secreted protein; CTSL is a protein located in the Golgi apparatus and additionally in vesicles; PAICS is a protein found in the cytosol while DAAM1 is located in the plasma membrane and cytosol (**Figure 1E**). Finally, we analyzed the gene expression profile of *ACE2, TMPRSS2, CTSL, DPP4, DAAM1*, and *PAICS* genes on publicly available scRNA-Seq datasets of lung, liver, and thymus fetal tissues. *CTSL*, *DPP4*, and *DAAM1* were found as highly expressed in fetal cells compared to *ACE2* and *TMPRSS2* in all tissues analyzed (**Supplementary Figure 2**). Additional investigations may determine the generality and impact of these findings, including the confirmation of vertical transmission.

**Figure 2.**
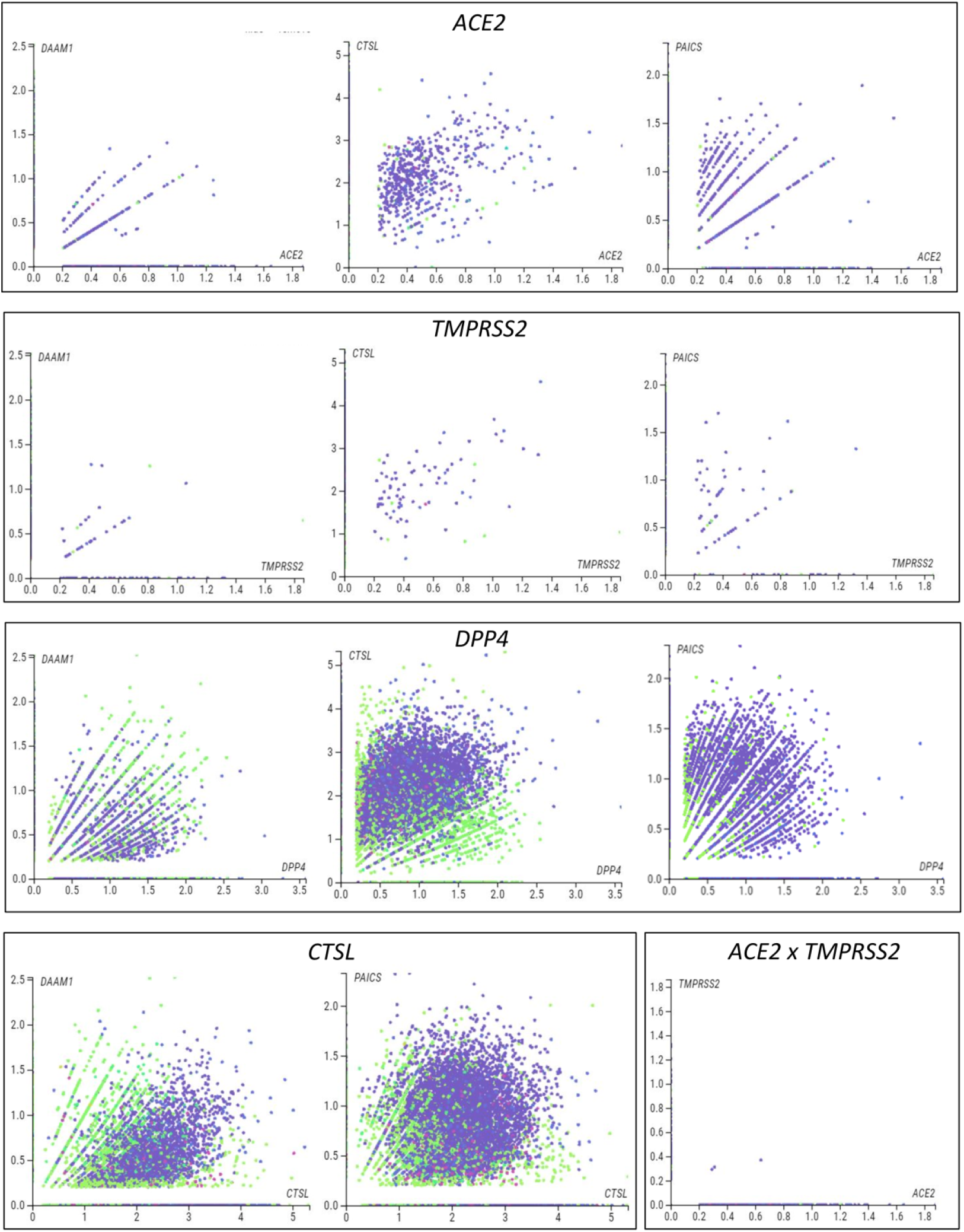
Single-cell analysis of the human placenta indicates potential non-canonical routes of SARS-CoV-2. **A**. Co-expression analysis of *DAAM1, CTSL*, DPP4, and *PAICS* with the canonical virus entry-associated genes *ACE2* and *TMPRSS2*. **B**. Co-expression analysis of *DAAM1* and *PAICS* with the non-canonical virus entry genes *DPP4* and *CTLS*. The images were generated using COVID-19 Cell Atlas (https://www.covid19cellatlas.org) functionalities.

## Discussion

Recent literature has shown placenta cells infected with SARS-CoV-2 in pregnant women diagnosed with moderate to severe COVID-19 (*15*, *22*) with findings supporting the possibility of vertical transmission of SARS-CoV-2 (*15*, *23*). However, few placenta cells express *ACE2* and *TMPRSS2* (*18*, *19*) required for virus entry, raising the question about whether potential non-canonical molecular mechanisms may be involved in the biological cycle of the virus in these cells. Here, we demonstrated by transcriptomic analyses (microarray and scRNA-Seq) and *in silico* predictions of virus-host protein-protein interactions that, despite low-levels of *ACE2* and *TMPRSS2*, villous trophoblast cells express high levels of the potential non-canonical cell-entry mediators *DPP4* and *CTSL*. We also found changes in the expression of *DAAM1* and *PAICS* genes that code for proteins predicted to interact with SARS-CoV proteins during pregnancy. These results provide evidence of potential host-virus PPI that may lead to SARS-CoV-2 infection and replication in the human placenta.

Firstly, we characterized the expression of SARS-CoV-2/SARS-CoV entry-associated receptors and proteases genes, which revealed that *DPP4* and *CTSL* are highly expressed in villous trophoblast cells during pregnancy. DPP4 has a high affinity to the SARS-CoV-2 spike receptor-binding domain, (*24*) and critical DPP4 residues share the SARS-CoV-2-S/DPP4 binding as found for MERS-CoV-S/DPP4 (*27*). In a pioneer study based on SARS-CoV cell entry mechanisms, Simmons et al. (*25*) described a three-step method for host-virus membrane fusion, involving receptor-binding and induced conformational changes in S glycoprotein with subsequent CTSL proteolysis and activation of membrane fusion within endosomes. During the COVID-19 pandemic, it has been further demonstrated that SARS-CoV-2 can use CTSB and CTSL as well as TMPRSS2 for priming host cells (*26*, *27*). These data provide evidence that CTSL is a potentially promising treatment for COVID-19 by blocking coronavirus host cell entry and intracellular replication (*28*). Secondly, we analyzed whether placental development and growth are associated with transcriptional changes in proteins that potentially interact with SARS-CoV. Among the transcripts translated into proteins predicted as interacting with SARS-CoV, *DAAM1* increased with placental development and growth. DAAM1 was previously identified as a regulator of bacteria phagocytosis by regulating filopodia formation and phagocytic uptake in primary human macrophages (*29*). We predicted that DAAM1 potentially interact with the viral protein encoded from open reading frame 8 (ORF8) and the hypothetical protein sars7a of SARS-CoV. The SARS-CoV ORF8 shares the lowest homology with all SARS-CoV-2 proteins; however, it was previously demonstrated that SARS-CoV-2 ORF8 expression was able to selectively target MHC-I for lysosomal degradation by an autophagy-dependent mechanism in different cell types (*30*). We found that *PAICS* transcript, which is also translated into a protein predicted as interacting with the Nsp3 of SARS-CoV, decreased its levels with placental development and growth. PAICS is an enzyme of de novo purine biosynthesis pathway, which was previously predicted as having a putative interaction with the human influenza A virus (IAV) nucleoprotein and was up-regulated during IAV infection (*31*). The putative interaction of Nsp3 with PAICS is relevant. Nsp3 is one of the 16 non-structural proteins in the replicase ORF1ab gene of SARS-CoV and SARS-CoV-2 (*1*, *32*), and binds to virus RNA and proteins, including the nucleocapsid protein (*33*). Considering the low-levels of *ACE2* and *TMPRSS2* that we found in villous trophoblast cells, our results provide evidence of alternatives cell-entry mediators in the placenta. Moreover, the description of the potential interaction between host and SARS-CoV-2 may provide insights into the mechanisms of placental infection in women with COVID-19.

Cellular and molecular composition play a role in placenta infection and intrauterine transmission of viruses (*34*, *35*). Thus, we finally used single-cell analysis of the syncytiotrophoblast, villous cytotrophoblast, and extravillous trophoblasts - the fetal component of the placenta - that confirmed the low-levels of expression of the canonical entry receptor *ACE2* and *TMPRSS2* genes (*18*, *19*). Conversely, we found that these cells express high-levels of *DPP4* and *CTSL*, two potential mediators of SARS-Cov-2 host cell entry (*24*, *27*) that may contribute to the SARS-CoV-2 infection (*15*, *22*, *23*). It is essential to highlight that our scRNA-Seq analysis showed that the non-canonical *DPP4* and *CTSL* entry genes (*24*, *27*) are highly co-expressed in syncytiotrophoblast, villous cytotrophoblast, and extravillous trophoblasts cells. The transcripts *DAAM1* and *PAICS*, which are translated into proteins predicted as potentially interacting with SARS-CoV-2, were also highly co-expressed with *DPP4* and *CTSL*. These results demonstrate that, although *ACE2* and *TMPRSS2* are poorly expressed in the placenta, other mediators that potentially interact with the virus are highly co-expressed in villous trophoblast cells and, therefore, may represent a valuable alternative route for infection and viral replication.

Although our *in-silico* analyses are a starting point, they have limitations. We reanalyzed published placenta datasets with a limited number of samples (microarray = 12 individuals, and scRNA-seq = 5 individuals) and, consequently, validations in a large cohort of samples must be performed. The predicted interactions presented here should also be experimentally validated in infected cells or placenta to circumvent these limitations. Moreover, the analyses of single-cell data should be conducted for the entire period of gestation, considering that we only evaluated single-cell data from the first trimester of pregnancy. The detection of SARS-CoV-2 in the placenta needs to be performed in a large cohort of pregnant women with COVID-19, including asymptomatics. Considering that vertical transmission of SARS-CoV-2 is still debated, the follow-up of newborns from mothers with COVID-19 during pregnancy should be necessary since, if it occurs even in the non-asymptomatic, the long-term consequences are mostly unknown.

In conclusion, despite low-levels of *ACE2* and *TMPRSS2*, our analyses demonstrate that villous trophoblast cells express high levels of potential non-canonical cell-entry mediators *DDP4* and *CTSL*. We also found changes in the expression of the *DAAM1* and *PAICS* genes coding for proteins predicted to interact with SARS-CoV proteins during pregnancy. These results provide new insight into the interaction between SARS-CoV-2 and the host proteins and indicate that coronaviruses may use multiple mediators for virus infection and replication.

## Materials and Methods

### Differential expression analysis of the placenta during pregnancy trimesters

We reanalyzed microarray data from villous trophoblast tissues of first (45-59 days, n = 4), second trimester (109-115 days, n = 4), and third trimesters (n = 4) available in Gene Expression Omnibus (GEO), under the accession number GSE9984 (*36*). Next, we identified the DEGs during the pregnancy, by comparing the second trimester and third trimester with the first trimester using the GEO2R tool (https://www.ncbi.nlm.nih.gov/geo/geo2r. Genes with Log2 of Fold Change ≥ |0.5| and False Discovery Rate (FDR) < 0.05 were considered as DEGs. We further grouped the gene expression profiles during pregnancy by using the K-means clustering analysis based on One Minus Pearson Correlation (Robust Multi-array Average, RMA, normalization log2, K-means = 3).

### Enrichment analysis

We used the Metascape tool (https://metascape.org) (*37*) to perform functional enrichment analysis of the genes lists generated in the clustering and differential expression analyses.

### Placenta proteins that potentially interact with coronaviruses and Zika virus

SARS-CoV-2 cell entry mediators were selected for first screening using the human proteins already described in the COVID-19 Cell Atlas (https://www.covid19cellatlas.org/) and the literature (*18*, *24*, *27*, *38*). Next, we used gene expression data to generated a list of PPIs that potentially interact with human coronaviruses and the ZIKV from the Pathogen–Host Interactome Prediction using Structure Similarity (P-HIPSTer, http://phipster.org/) database (*23*) (**Supplementary Table 4 and 5, respectively**). SARS-CoV was included in the analysis considering the evolutionary relationship between the novel SARS-CoV-2 (*38*), and ZIKV considering its impact on placental infection and fetal microcephaly (*7*, *34*).

### Tissue-specific gene networks of potential virus-host placenta interactome

The HumanBase webtool (https://hb.flatironinstitute.org) (*39*) was used to generate the tissue-specific gene network of placental genes coding for proteins that potentially interact with SARS-CoV-2 based on P-HIPSTER or that we identified as either associated with COVID-19 or that we hypothesized may be associated with the disease, based on the literature (**Supplementary Table 8**). We considered co-expression, interaction, transcriptional factor binding, Gene Set Enrichment Analysis (GSEA) perturbations as active interaction sources, applying minimum interaction confidence of 0.07, and a maximum number of 15 genes.

### Single-cell transcriptome analysis of human placentas and fetal tissues

We used the functionalities of the COVID-19 Cell Atlas (https://www.covid19cellatlas.org) to explore single-cell RNA sequencing (scRNA-Seq) data from five samples from first-trimester placentas, previously described by Vento-Tormo et al., (*40*). The expression of placental genes potentially interacting with SARS-CoVs was visualized in placental cell types using the “interactive viewer” developed by cellxgene v0.15.0 (https://cellxgene.cziscience.com). We also characterized the gene expression profile of canonical and non-canonical placental mediators of viruses interaction per cell using cellxgene v0.15.0 bar plots. The placental single-cell transcriptome is available for re-analysis at Human Cell Atlas Data Portal (https://data.humancellatlas.org) or E-MTAB-6701 (EMBL-EBI, ArrayExpress; https://www.ebi.ac.uk/arrayexpress). We applied this same strategy to investigate the gene expression of our candidates using scRNA-seq in human fetal tissues: lung, liver, and thymus (ArrayExpress; E-MTAB-8221, E-MTAB-7407, and E-MTAB-8581, respectively. Last accessed June 2020.

### Data representation and analysis

Morpheus (https://software.broadinstitute.org/morpheus/) was used to generate the heatmaps and to perform the clustering analyses. The Venn diagram was constructed using the Van de Peer Lab software (http://bioinformatics.psb.ugent.be/webtools/Venn/). Biorender was used to design cell images (https://app.biorender.com/).

### Data Availability Statement

All data is available in the manuscript.

## Author contributions

Conceptualization, F.B.C., R.F.C., L.A.J.; methodology, F.B.C., R.F.C., S.S.C., L.A.J.; formal analysis, F.B.C., C.R.N., R.F.C., S.S.C., L.A.J; investigation, F.B.C., C.R.N., R.F.C., S.S.C., L.A.J.; resources, R.F.C., L.A.J.; data curation, F.B.C., C.R.N., R.F.C., S.S.C., L.A.J.; writing—original draft preparation, F.B.C., R.F.C., S.S.C., L.A.J.; writing-review and editing, all authors; supervision, R.F.C., L.A.J; project administration, R.F.C., L.A.J; funding acquisition, R.F.C., L.A.J.

## Ethical Approval

Not applicable as we used publicly available data.

## Supplementary Figures

**Supplementary Figure 1.**
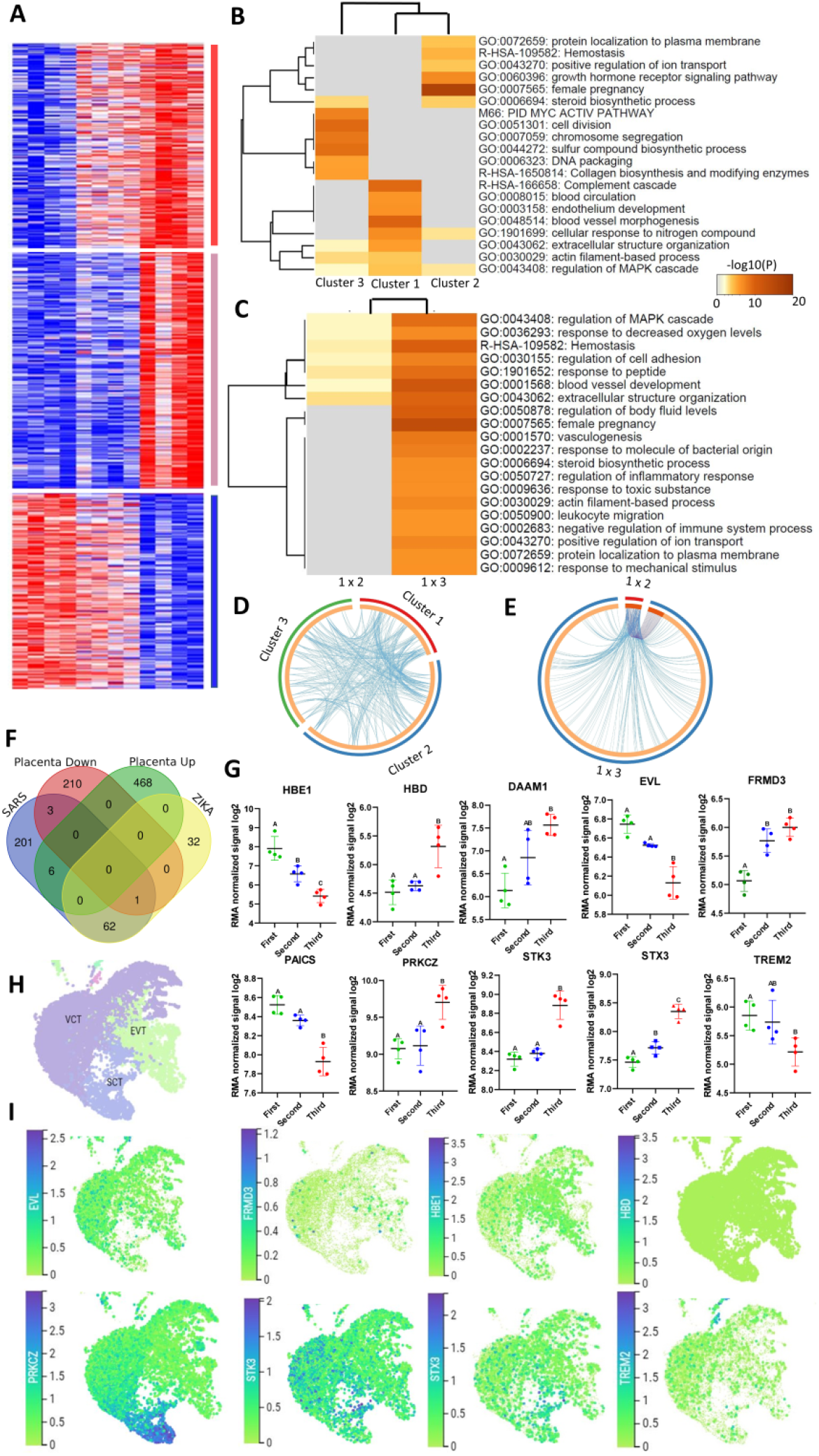
Gene expression profile of gestational trimesters and its potential to interact with SARS-CoV-2 proteins. **A**. Heatmap illustrates the expression (counts RMA normalized) of placenta differentially expressed genes (DEGs) from the three trimesters (GSE9984). Clustering analysis was performed based on One Minus Pearson Correlation and k-means analysis generated three clusters related to the increase or decrease of the gene expression during the gestation. Cluster 1 is represented by the red bar, cluster 2 by the pink bar, and cluster 3 by the blue bar. **B**. Functional enrichment analysis generated in the Metascape tool (https://metascape.org) (*37*) from each cluster found in the placenta DEGs. **C**. Functional enrichment analysis generated in the Metascape tool from DEGs belonging to the comparison of first versus second trimester and first versus term placentas. **D**. Circus plot showing overlapped functional enriched terms from the gene lists of the clusters (1, 2 and 3). **E**. Circus plot showing the overlapping between the functional enriched terms generated from DEGs of first versus second trimester and DEGs from first trimester versus term placentas. **F**. Venn Diagram showing the DEGs shared by the placenta (up and down), human protein of expected interaction with SARS-CoV, and human protein of expected interaction with ZIKA virus. **G**. Box plot showing the expression of the 10 DEGs encoding for proteins with expected interaction with SARS-CoV. **H-I**. Single-cell gene expression analyses in placental cells types using scRNA-Seq data from Vento-Tormo et al, 20181. Dimensionality reduction demonstrates different cell populations identified in five human placental samples in a Uniform Manifold Approximation and Projection (UMAP) plot. SCT, syncytiotrophoblast; VCT, villous cytotrophoblast; EVT, extravillous trophoblast. The images were generated using COVID-19 Cell Atlas (https://www.covid19cellatlas.org) functionalities. RMA: Robust Multichip Average.

**Supplementary Figure 2.**
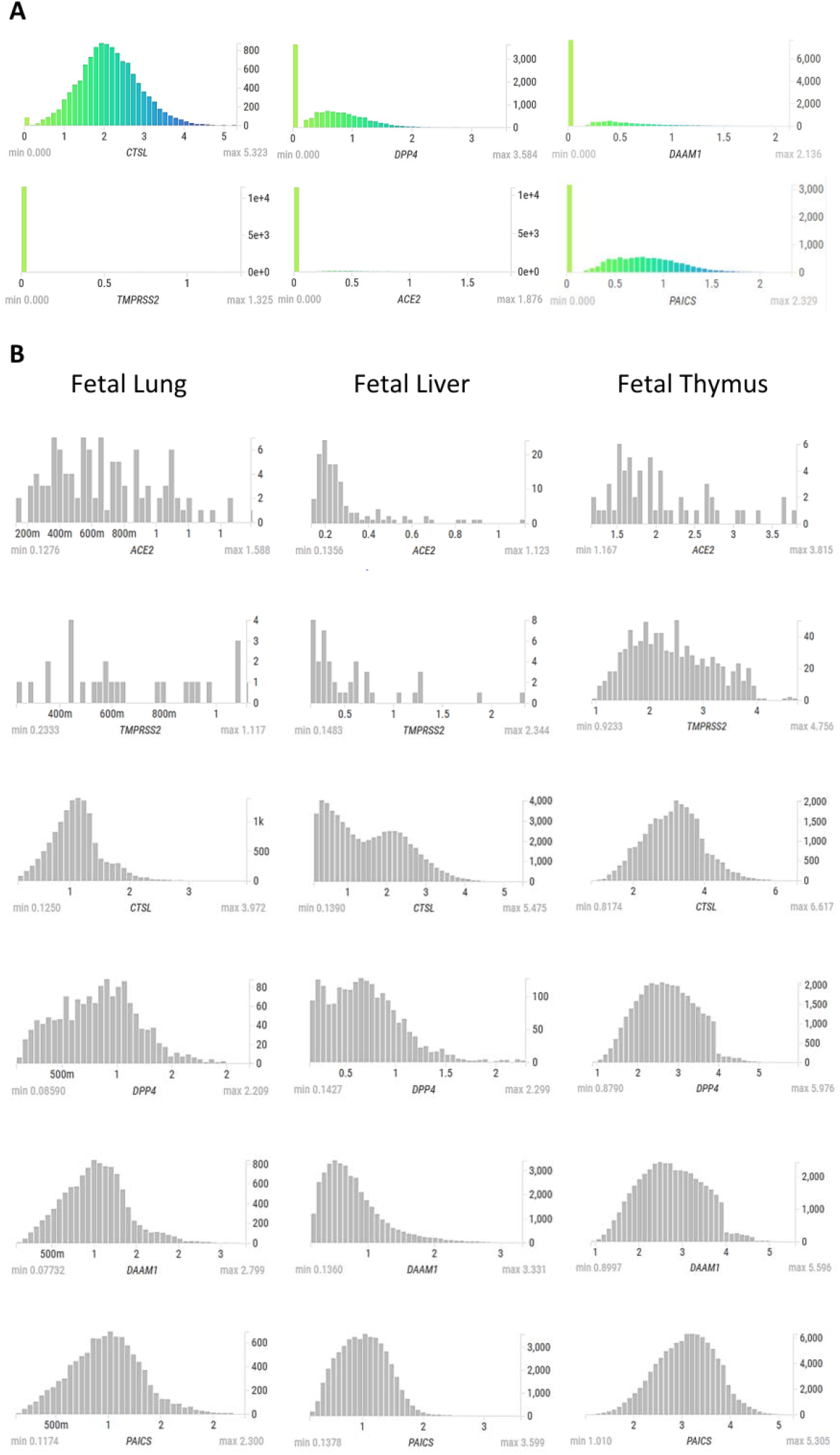
Gene expression of ACE2, TMPRSS2, CTSL, DPP4, DAAM1, and PAICS in scRNA-seq of fetal tissues. **A**. Bar plots representing the gene expression (log-transformed normalized counts) of *ACE2, TMPRSS2, CTSL, DPP4, DAAM1*, and *PAICS* in human placental tissues (Vento-Tormo et al., 2018). X-axis indicates log-transformed normalized counts while Y-axis indicates the amount of cell expressing the gene. **B**. Bar plots representing the gene expression (log-transformed normalized counts) of *ACE2, TMPRSS2, CTSL, DPP4, DAAM1*, and *PAICS* in human fetal lung (Miller et al., 2020), liver (Popescu et al., 2019), and thymus (Park et al., 2020). X-axis indicates log-transformed normalized counts while Y-axis indicates the amount of cell expressing the gene. The images were generated using COVID-19 Cell Atlas (https://www.covid19cellatlas.org) functionalities and the cells not expressing counts were omitted from the plot.

## Supplementary Tables S1-S8

https://drive.google.com/file/d/1qUIFjAc28Dk0TIPDCURR59ewIamgkBv6/view?usp=sharing

**Supplementary Table 1**. Differentially expressed genes in placenta tissue during the gestation (1° trimester x 2° trimester and 1° trimester x 2° trimester; Log2FC >= |0.5|; FDR< 0.05).

**Supplementary Table 2**. Functional enrichment analysis generated in the Metascape tool (https://metascape.org) (*37*) for each cluster generate for placenta gene expression during gestation.

**Supplementary Table 3**. Functional enrichment analysis list generated in the Metascape tool (https://metascape.org) [DOI: 10.1038/s41467-019-09234-6] for DEGs in the placenta tissues (1° trimester x 2° trimester and 1° trimester x 2° trimester.

**Supplementary Table 4**. Human-SARS-CoV Interactome based on the in silico computational framework P-HIPSTer (http://phipster.org/).

**Supplementary Table 5**. Human-ZIKA Interactome based on the in silico computational framework P-HIPSTer (http://phipster.org/).

**Supplementary Table 6**. The Human-SARS-CoV interactome obtained in the P-HIPSTer (http://phipster.org/) for placenta.

**Supplementary Table 7**. The Human-ZIKA interactome obtained in the P-HIPSTer (http://phipster.org/) for placenta.

**Supplementary table 8**. Human-SARs Interactome based on literature.

