## Supplementary figures and images for "Prediction of non-canonical routes for SARS-CoV-2 infection in human placenta cells"

### Supplementary figure 1

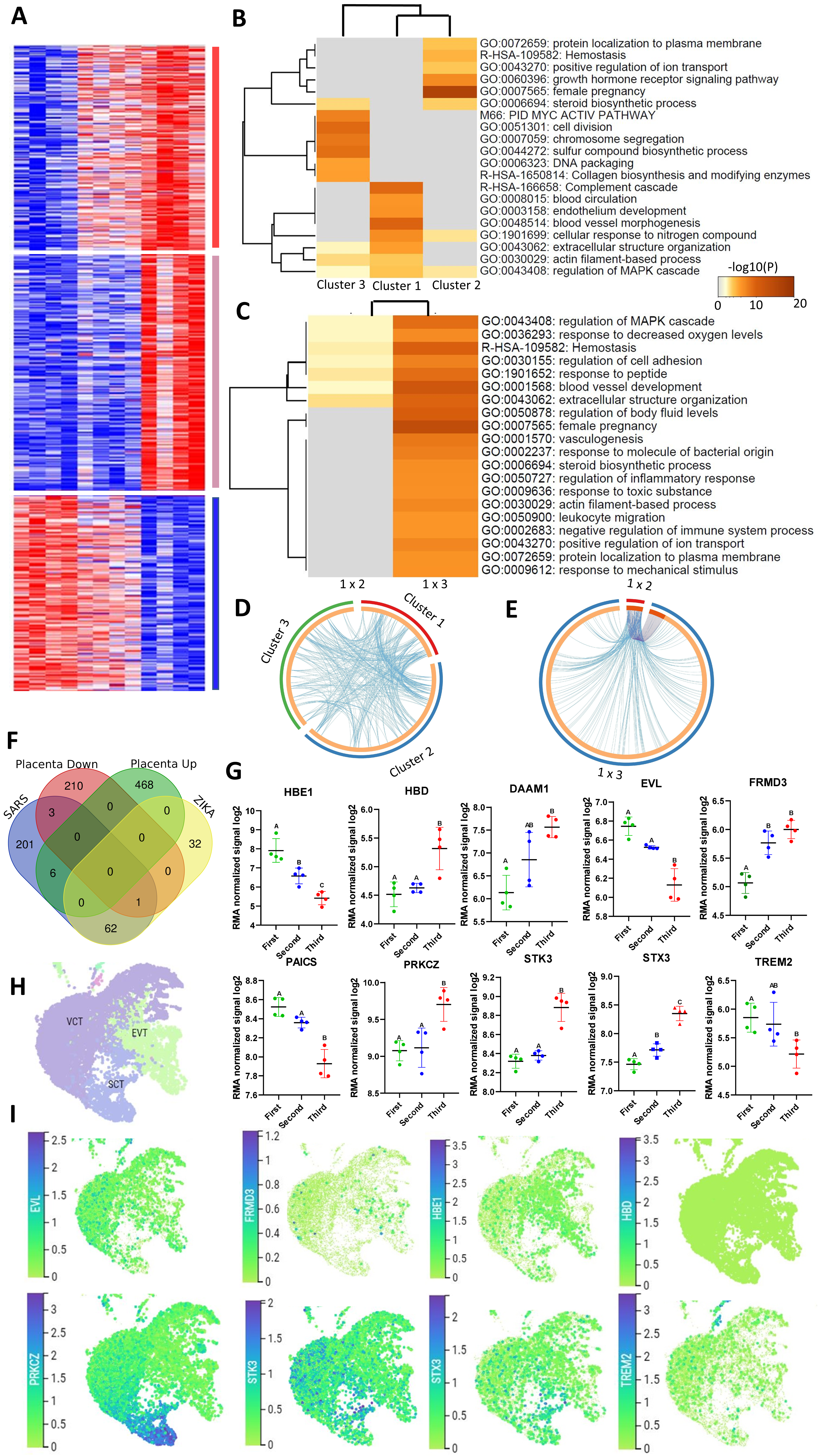

### Supplementary figure 2

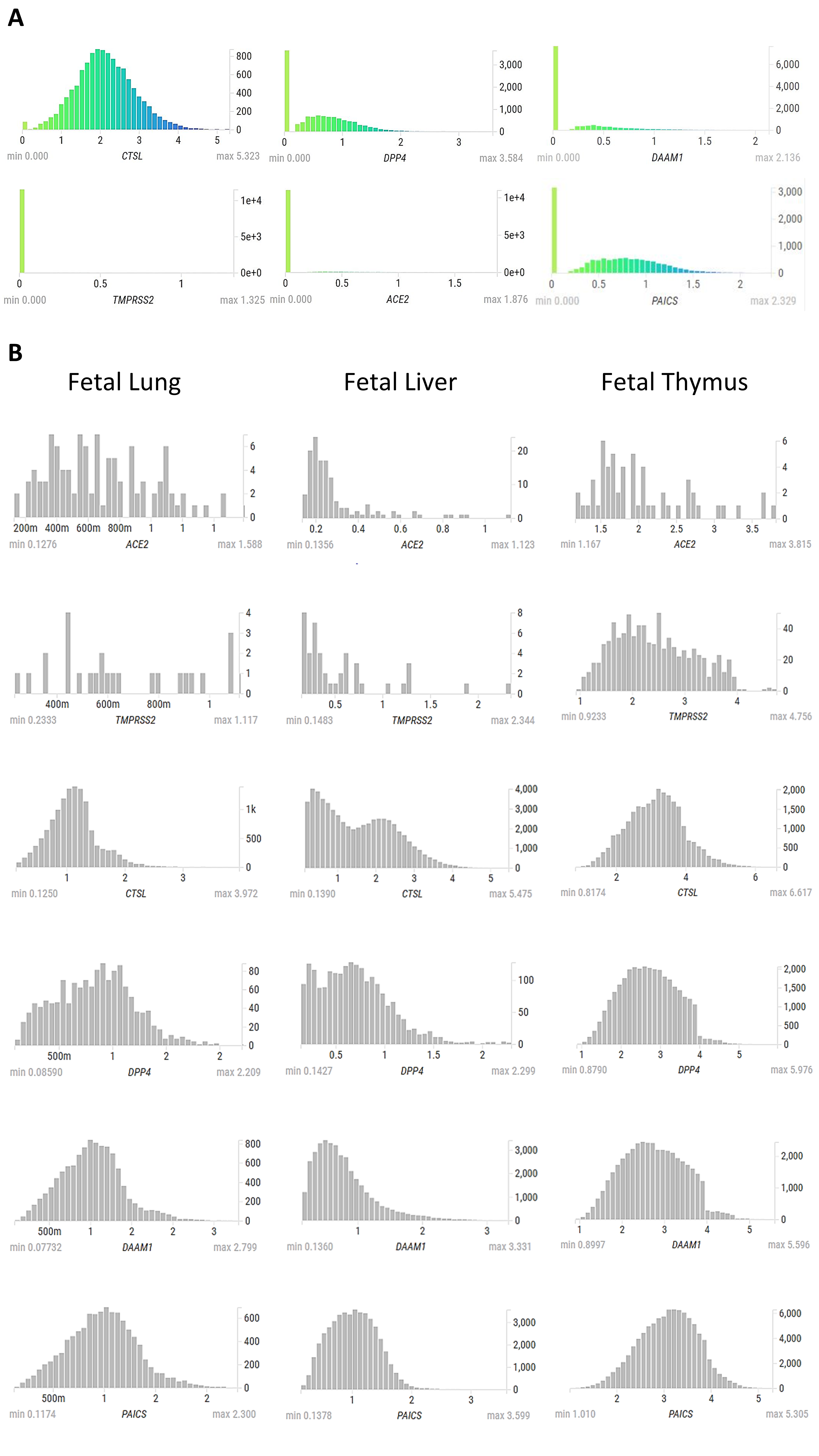
